# Modeling Genetic Diversity in Sickle Cell Disease Reveals Heterogeneous Responses to HbF-Inducing Therapies

**DOI:** 10.64898/2026.05.18.726003

**Authors:** Braden Pate, Anna Goldstein, Madeline Labott, Maria Lizarralde-Iragorri, Alisa Chankhunthod, Tyonna Tyson, Meghan Sloan, Charith Wijeyesekera, Andrew Wilks, Martin H Steinberg, George J Murphy, Kim Vanuytsel

## Abstract

Sickle cell disease (SCD) is caused by a point mutation in the β-globin gene that promotes hemoglobin polymerization, leading to chronic hemolytic anemia, vaso-occlusive episodes, and progressive organ damage. The most efficacious therapies focus on reactivating fetal hemoglobin (HbF) expression to mitigate the pathological effects of sickle hemoglobin (HbS) polymerization. However, the predominantly used HbF inducer, hydroxyurea (HU), exhibits substantial interpatient variability in efficacy, and curative approaches such as gene therapy remain inaccessible to the vast majority of patients. Although all SCD patients share the same causative HBB glu7val mutation, differences in genetic background significantly influence disease severity and therapeutic response. We describe a SCD-specific induced pluripotent stem cell (iPSC) platform as a renewable and scalable preclinical model to interrogate treatment responses across the genetically diverse SCD patient population. By generating patient-specific iPSC-derived erythroblasts (iEry) representing distinct SCD genetic backgrounds, we demonstrate that this system faithfully recapitulates the heterogeneous HbF induction observed clinically in response to HU. Moreover, this platform enables the identification and evaluation of alternative therapeutic agents for HU non-responders and provides sufficient resolution to dissect drug-specific effects on erythroid differentiation and cellular phenotypes. Together, these findings support the use of iPSC-derived erythroid models as a versatile tool to advance precision therapeutic strategies for SCD.

**KEY POINTS:**

- SCD iPSC-derived erythroid cells (iEry) reflect the diversity in HU-mediated HbF induction seen in SCD patients
- SCD iEry recapitulate patient-specific treatment responses and can be used to identify therapeutic alternatives for HU non-responders
- iEry provide a versatile platform to study the impact of novel HbF inducers on erythroid cell characteristics and differentiation parameters

## INTRODUCTION

Sickle cell disease (SCD) results from an A to T transversion at position 7 of the β-globin gene (*HBB*) that renders the mutant sickle hemoglobin (HbS; α_2_β^S^_2_) prone to polymerization. This aberrant polymerization is central to the pathophysiology of the disease, triggering many downstream events including hemolytic anemia, acute vaso-occlusive episodes, and chronic organ damage, severely impacting the span and quality of life^1,2^. Fetal hemoglobin (HbF; α_2_γ_2_), the predominant hemoglobin during fetal development that is replaced by adult hemoglobin (HbA; α_2_β^A^_2_) shortly after birth, interferes with HbS polymerization^3,4^. Therefore, one of the main therapeutic strategies is re-activation of HbF. Hydroxyurea (HU), the sole approved pharmacologic HbF inducer, is recommended for nearly all patients^5^, however the response is variable^6-9^. Curative cellular therapeutics like allogeneic transplantation and gene therapy have barriers to access in terms of cost, eligibility and availability^10-17^. As such, there is a continued need for investment in HbF-inducing drugs that can manage the disease in resource-limited areas of the world and for those not eligible for cell therapies.

While SCD patients all carry the same point mutation in their β-globin gene, differences in genetic context drive the diversity observed in disease severity and treatment response^9,18^. Assessing the therapeutic effects of novel compounds across a broad range of primary patient samples to account for this heterogeneity is impractical due to limited sample availability, experimental variation between samples, and the finite amount of material in each specimen. Alternatively, HbF induction studies widely use immortalized erythroid progenitor lines like HUDEP-2^19,20^ and BEL-A^21,22^. To increase disease relevance, the BEL-A cell line has been genetically engineered to carry disease-causing mutations, generating isogenic cell lines reflecting a SCD and β-thalassemia context^23^. Although these advances expand the investigative toolbox, each model represents only a single genetic background. Notably, some established HbF-inducing drugs show little effect in the commonly used HUDEP-2 cell line^22,24^. Relying on a limited number of backgrounds to evaluate a compound may therefore misrepresent its true clinical efficacy across a diverse patient population. The risk of a novel drug candidate failing in patients after significant investment highlights the need for more predictive preclinical models earlier in the drug development pipeline^25^

We sought to improve the range of genetic diversity that can be considered in a preclinical setting by relying on the ability of induced pluripotent stem cells (iPSCs) to capture the exact genetic background of a patient. iPSCs can be derived from somatic patient material and offer an unlimited source of cells that can be specified into any cell type, including red blood cells. We previously established a comprehensive, ethnically diverse SCD iPSC library that can be used to generate patient-specific erythroid cells *in vitro*^26^. We have shown that these cells can be efficiently differentiated towards the erythroid lineage^27^, and have extensively characterized the developmental and maturational status of iPSC-derived erythroid (iEry) cells^28,29^. Using a β-globin reporter iPSC line to track globin expression at the single cell level, we were able to demonstrate that iEry generated using our erythroid differentiation protocol represent definitive erythroid cells^28^. Immunophenotypic, transcriptomic and proteomic profiling, confirmed that iEry are in many ways similar to primary postnatal red blood cells^28,30^. One major difference between iEry and their postnatal counterparts is their predominant HbF expression, reflecting a prenatal stage of development prior to globin switching. Importantly, we show that the predominant expression of HbF in iEry, is not an obstacle for HbF induction screening. Here, we highlight the potential of this SCD iPSC platform as a versatile preclinical screening tool to assess HbF induction across a diverse SCD patient population. By generating patient-specific erythroid cells representing different SCD genetic backgrounds and *HBB* haplotypes, we show 1) a differential response to HbF-inducing agents across SCD patient backgrounds, 2) the utility of this platform to identify alternatives for patients who do not respond to commonly used treatments like HU, and 3) the resolution this renewable cell source offers in terms of drug-specific impact on erythroid cell characteristics and differentiation.

## METHODS

### iPSC generation and maintenance

iPSC lines (BU6, SA-170, SS-100, BR-SP-31, SS-24, SS-8) were created using previously established methodologies and met stringent quality control parameters for pluripotency and functionality^26,31,32^. All lines were maintained with mTeSR 1 (StemCell Technologies; #85850) and mTeSR Plus (StemCell Technologies; #100-0276) on matrigel (Corning Matrigel hESC-qualified Matrix; #354277) and passaged approximately every 7 days using ReLeSR (StemCell Technologies; #100-0484). All cells were cultured at 37ºC in normoxic, 5% CO2 conditions.

### Erythroid differentiation of iPSCs

iPSC cells were plated on matrigel (Corning Matrigel hESC-qualified Matrix; #356234) at ∼10% confluency and in mTeSR 48 hours prior to differentiation. To specify iPSC towards HSPCs, we followed a step-wise protocol as previously described^28,29^, with the following changes: For day 0 and 1, RPMI-1640 and 10% KO serum replacement were substituted with StemPro-34 SFM, and CHIR99021 was used on these days in place of hWnt3a. On day 2, the media previously described for day 3 was used. 10µM of CH223191 (Tocris; #3858) was added to day 4 and 5 media. The concentration of recombinant human SCF (hSCF) in the HSPC media was doubled to 200ng/ml.

On day 13, suspended HSPCs were collected and resuspended in SS1 media for further specification towards the erythroid lineage. SS1 media consists of the following: StemSpan SFEM II (StemCell Technologies; #09655), 2mM l-glutamine (Gibco; #25030164), 100ug/mL Primocin (InvivoGen; #ant-pm-2), 100ng/ml hSCF (R&D Systems; #255-SC), 40ng/mL recombinant human IGF-1 protein (IGF-1) (R&D Systems; 291-G1), 0.5μM Dexamethasone (Sigma-Aldrich; #D4902), 0.5 U/ml human erythropoietin (EPO) (Henry Schein; #1265340). iEry were collected on day 21. All cells were cultured at 37ºC in normoxic, 5% CO2 conditions.

### HbF induction testing

Cells were treated with Hydroxyurea (HU) (Sigma-Aldrich; #H8627), pomalidomide (POM) (Sigma-Aldrich; #P0018), decitabine (DAC) (Sigma-Aldrich; #A3656) or SR-19292 (Cayman Chemical; #22084) shortly following erythroid specification on days 15, 18, and 20. Cells were treated with RN-1 (Sigma-Aldrich; #489479) on days 18 and 20.

Compounds were prepared as per the manufacturer’s recommendations. Each assay included vehicle controls (H2O and DMSO).

### Wright-Giemsa staining

Cells were collected and made adherent to glass slides using cytospin centrifugation. Wright-Giemsa Staining was performed as directed in the Hema 3 Stain Kit Protocol (Fisher Scientific; #23-123869).

### Isolation, expansion and differentiation of primary CD34+ cells

Peripheral blood was collected in Vacutainer CPT Tubes (BD; #362760) and centrifuged for 30min at 1800 x G at room temperature (RT). The buffy coat was extracted and washed for 15min at 300 x G at RT using phosphate buffer saline (PBS) to recover the peripheral blood mononuclear cell (PBMC) cell pellet. CD34+ cells were enriched using the EasySep Human CD34 Positive Selection Kit II (StemCell Technologies; #17856) and EasySep Magnet (StemCell Technologies; #18001) according to the manufacturer’s recommendations.

Prior to differentiation, CD34+ cells were expanded for 5-7 days in HSC Expansion Media consisting of: StemSpan SFEM II, 100ng/ml hSCF, 100ng/ml Recombinant Human Flt-3 Ligand (FLT3L) (R&D Systems; #308-FK), 10ug/ml Recombinant Human Thrombopoietin Protein (hTPO) (R&D Systems; #288-TP), and 100ng/ml Human low-density lipoproteins (hLDL) (StemCell Technologies; #02698). Subsequent erythroid differentiation was performed according to previously described methodologies^33^. HU was added to the medium starting on day 2 of culture, and replenished with each addition of fresh medium, for a read-out of HbF induction on days 7 (shown in Figure 1), 11 and 13. All cells were cultured at 37ºC in normoxic, 5% CO2 conditions.

**Figure 1.**
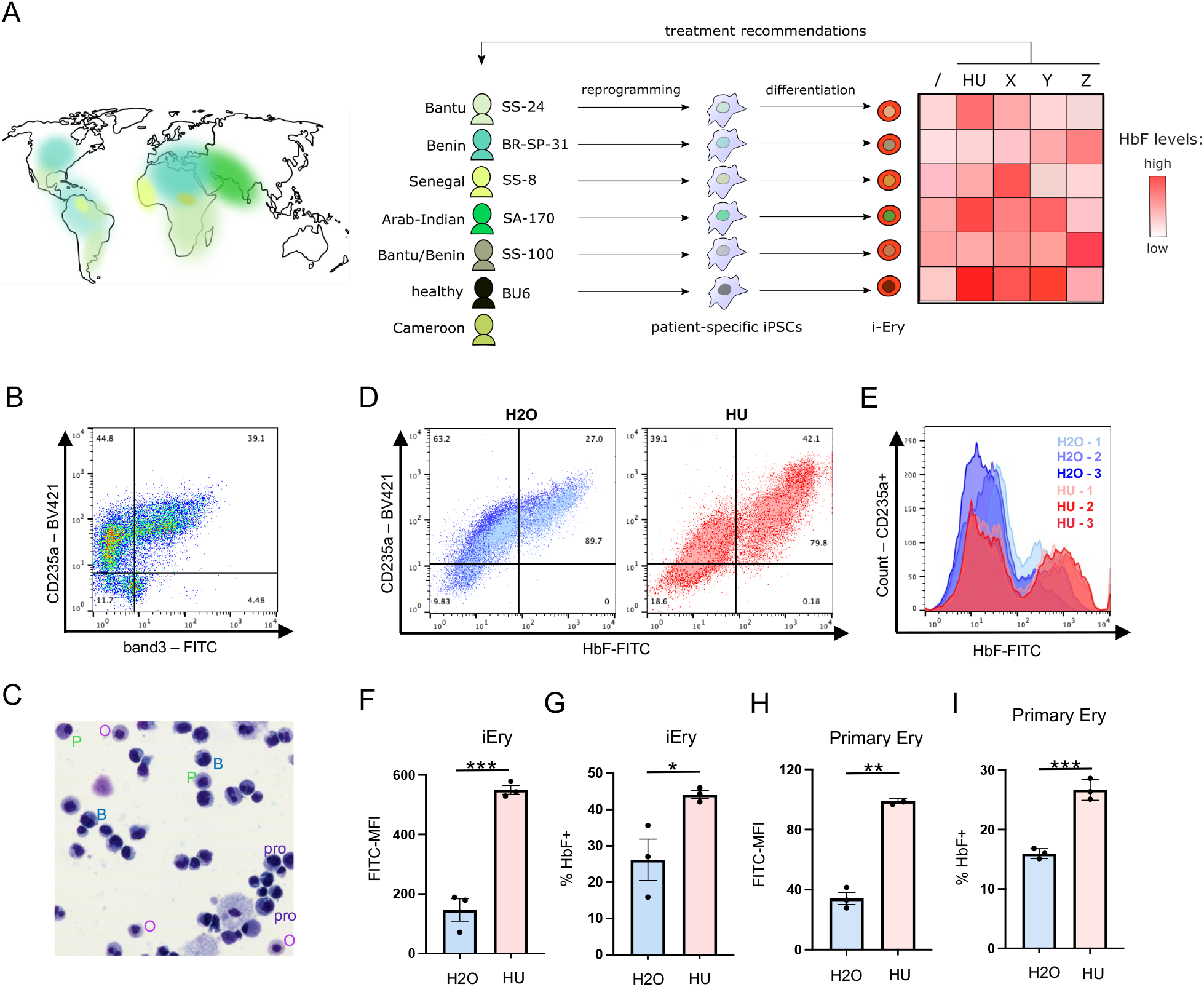
Baseline HbF expression in iEry is not an obstacle to HbF induction screening. A) Conceptual overview: Distribution of the HbS gene with different colors representing different *HBB* haplotypes and their areas of greatest density. Representative iPSC lines corresponding to haplotypes used in this study are indicated. Heatmap depicts hypothetical outcomes in terms of HbF induction across treatments (HU, X, Y, Z) and backgrounds. B) Representative flow cytometry plot of day 21 erythroid cultures (BU6). C) Wright-Giemsa staining of day 21 erythroid cultures (BU6): proerythroblast (pro), basophilic (B), polychromatic (P) and orthochromatic (O) erythroblasts. D) Flow cytometry plots illustrating HbF induction following 6 days of HU [25µM] treatment (BU6, n=2). E) HbF induction following 6 days of HU [25µM ] treatment represented as histograms (BU6, n=2). F) Quantification of HbF induction via Mean Fluorescence Intensity (MFI) of HbF-FITC for HU [25µM] and H_2_O conditions (BU6 iEry, n=2). G) %HbF+ cells as measured by flow cytometry (BU6 iEry, n=2) H) Quantification of HbF induction via Mean Fluorescence Intensity (MFI) of HbF-FITC for HU [25µM] and H_2_O conditions (BU6 primary erythroid cells, n=3) I) %HbF+ cells as measured by flow cytometry (BU6 iEry, n=3). Statistical significance between experimental groups was determined using a Student’s t test. A p value <0.05 was considered significant: *p < 0.05 and ***p < 0.001. Error bars represent mean ± standard error of mean (SEM)

### FACS Analysis

Cells were stained on ice for 30 minutes using the following antibodies: CD34-BV421 (BD; #562577), CD45-PE (BD; #555483), CD235a-BV421 (BD; #562938), Band3 (CD233)-FITC (American Research Products; #08-9439-3), and HbF1-FITC (Invitrogen; #MHFH01). For combinations with HbF antibodies, fixation and permeabilization was performed prior to staining as directed in the Cytofix/Cytoperm Fixation/Permeabilization Kit (BD; #554714). Cells were read out on a Stratedigm S1000EXI and data was analyzed using FlowJo (v10) (FlowJo, LLC) software. Live cells were gated based on forward scatter and side scatter properties. Quantification of mean fluorescence intensity (MFI) and % HbF+ cells was performed on CD235a+ gated cells. HUDEP2 cells, expressing minimal to no HbF, were used as gating controls to determine the HbF+ gate.

### Statistics

Individual differentiations were conducted with 3 technical replicates (n=3) for each treatment condition, and for each cell line. When compiling data across differentiations (n=4-12), we used the average values from each differentiation for the conditions tested. Relative values (compared to vehicle control conditions H2O and DMSO) were calculated to account for variation in fluorescence intensity between read-outs.

Statistical analyses were performed using GraphPad Prism (v10.6). To determine significance, a one-sample t-test was performed on the fold-change values of the treatment group against a theoretical mean of 1.0 (control). Normality of the treatment group was confirmed via a Shapiro-Wilk test. For analyses where not all data was normally distributed, we applied a Wilcoxon test. When working with absolute values, an unpaired t-test was used to determine significance. A paired t-test was used for linked outcomes. When considering multiple treatments versus a control condition, we used an ordinary one-way ANOVA test with Dunnett correction.

### Haplotyping

Haplotyping of SA-170, SS-8, SS-24 and BR-SP-31 lines was performed previously^26^. To determine the haplotype of SS-100, restriction fragment length polymorphism (RFLP) haplotype analysis was performed via digestion of the following polymerase chain reaction (PCR)-amplified polymorphic restriction sites within the beta globin gene cluster: Hinc II 5’ to ε, XmnI 5’ to G γ, Hind III in G γ, and Hinc II 5’ to δ (Figure S1A). DNA was isolated using the DNeasy Blood & Tissue Kit (Qiagen; #69506) and restriction site-containing DNA sequences were amplified using 2x GoTaq Green Master Mix (Fisher; #M7123), using primers (Integrated DNA Technologies) based on previously published sequences (see Supplemental Table 1). Restriction enzymes and buffers were all purchased from New England Biolabs.

## RESULTS

### SCD iEry reflect the diversity in HU-mediated HbF induction observed in the patient population

To assess HbF induction capacity across patient genetic backgrounds, we selected SCD iPSC lines representing various *HBB* haplotypes to capture the diversity of the patient population (**Fig. 1A**). Using a directed differentiation protocol designed to specify iPSC towards definitive erythroid cells^28,29^, we differentiated six iPSC lines: one healthy control (BU6) and five SCD patient lines corresponding to Bantu (SS-24), Benin (BR-SP-31), Arab-Indian (SA-170), Senegal/Cameroon (SS-8) and Bantu/Benin (SS-100) haplotypes^26^ (**Fig. 2A, Fig. S1A**). Similar to our previous report^26^, these lines all efficiently generated CD34^+^ CD45^+^ HSPCs and subsequently CD235a-expressing erythroid progeny (**Fig. 1B and S1B-C**).

**Figure 2.**
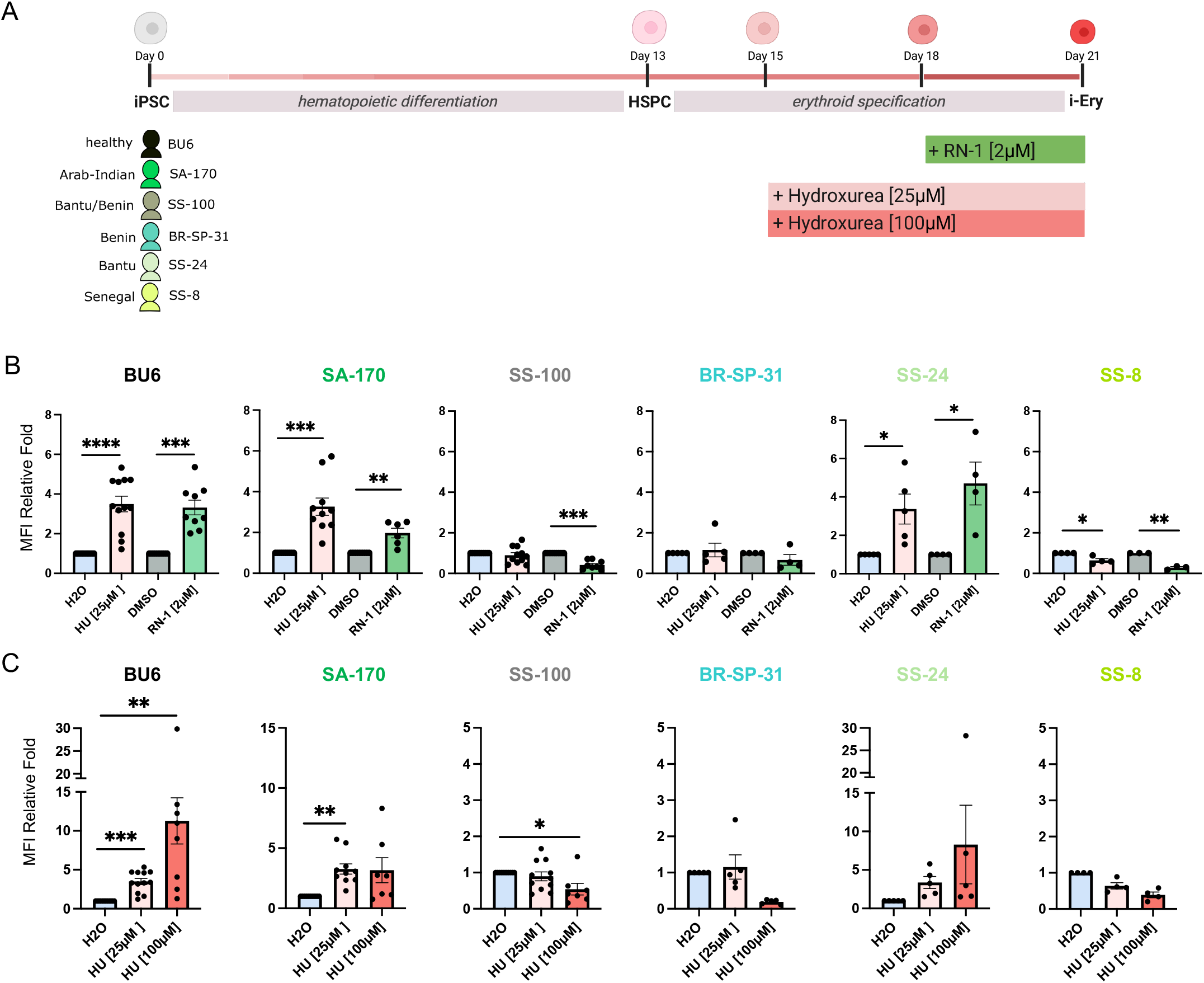
SCD iEry reflect the diversity in HU-mediated HbF induction observed in the SCD patient population. A) Schematic overview of differentiation and dosing process. B) HbF induction following HU and RN-1 treatment quantified by Mean Fluorescence Intensity (MFI) relative to control conditions H2O and DMSO (n=4-12 independent differentiation and dosing experiments, considering the average relative fold MFI of 2-3 repeats per differentiation). C) HbF induction following different HU doses quantified by Mean Fluorescence Intensity (MFI) relative to control condition H2O (n=4-12 independent differentiation and dosing experiments, considering the average relative fold MFI of 2-3 repeats per differentiation). Statistical significance between experimental groups was determined using a one-sample t-test (B) or Wilcoxon test (C). A p value <0.05 was considered significant: *p < 0.05, **p < 0.05, ***p < 0.001 and ****p < 0.0001. Error bars represent mean ± standard error of mean (SEM)

To validate the suitability of our iPSC-derived cells to report on HbF induction, we first assessed the capacity of healthy control BU6 iEry to induce HbF expression upon HU treatment. Shortly after specification towards the erythroid lineage (day 15), cells were dosed with HU (25µM) and changes in HbF expression were analyzed by flow cytometry 6 days later (day 21) (**Fig. 2A**), when cultures comprised erythroid progenitors spanning proerythroblast, basophilic, polychromatic and orthochromatic erythroblast stages as judged by flow cytometry and Wright-Giemsa staining (**Fig. 1B-C, Fig. S1C**). We found that HU treatment resulted in a 3.8-fold increase in mean fluorescence intensity (MFI) of the HbF-FITC signal (**Fig. 1D-F**) as well as a 1.7-fold increase in % HbF^+^ cells compared to iEry treated with H2O (**Fig. 1D,G**). This was comparable to the HbF induction capacity of primary erythroid progenitors (predominantly expressing adult hemoglobin (HbA)) from the same BU6 background, where upon HU dosing a 2.9-fold increase in HbF-FITC MFI and a 1.7-fold increase in HbF^+^ cells was noted (**Fig. 1H,I**). Together these results illustrate that baseline HbF expression in iEry is not an obstacle to HbF induction screening.

We next assessed HbF induction capacity across the selected SCD backgrounds following dosing with HbF-inducing compounds HU and the lysine-specific demethylase 1 (LSD1)-inhibitor RN-1^34,35^ (**Fig. 2A**). Comparing HbF-FITC MFI between dosed and vehicle control conditions revealed significant differences in HbF induction across genetic backgrounds for both HU and RN-1 (**Fig. 2B**). More specifically, SA-170 and SS-24 iEry showed a more than 3-fold increase (3.3-fold and 3.4-fold respectively) in HbF-FITC MFI upon HU treatment, like the 3.5-fold increase in MFI seen in healthy control cells (BU6). The same patient backgrounds (SA-170 and SS-24), as well as BU6, showed an increase in HbF-FITC MFI upon RN-1 dosing of 2-, 4.7- and 3.3-fold respectively. iEry from other SCD backgrounds tested (SS-100, BR-SP-31 and SS-8) failed to demonstrate HbF induction in response to either treatment. These results reflect clinical observations, where some SCD patients show an increase in HbF expression upon HU treatment, whereas others do not.

To determine whether this lack of response was due to insufficient dosing, we repeated these experiments with a higher dose of HU (100µM) in parallel to the more standard dose of 25µM^36,37^ (**Fig. 2C**). For the healthy control line BU6, HU increased HbF expression at both 25µM (3.5-fold) and 100µM (11.3-fold) concentrations. We did not observe significant improvements in HbF induction capacity at 100µM for other backgrounds. SCD lines that did not show increased HbF expression at 25µM HU (SS-100, BR-SP-31 and SS-8) failed to show HbF induction even at 100µM HU, indicating that the absence of a response is not due to inadequate dosing.

### SCD iEry recapitulate patient-specific treatment responses and suggest therapeutic alternatives

Interestingly, the absence of HbF induction in SS-100 iEry following HU treatment (**Fig. 2B,C**) faithfully recapitulates the clinical phenotype of the donor. Despite years of HU therapy at maximum tolerated dose (1500-2000 mg), this subject exhibited little HbF response to HU and continued to experience >3 severe vaso-occlusive episodes per year. Leveraging our platform’s capacity for parallel testing across a renewable, patient-specific supply of cells, we evaluated alternative HbF-inducing agents for this HU-refractory background (**Fig. 3A**). In healthy control iEry (BU6), pomalidomide^38^ and decitabine^39,40^ treatment resulted in increased HbF expression (2.2-fold and 3.7-fold respectively), consistent with their effects reported in the literature and similar to the effect noted for HU and RN-1 in this background (**Fig. 3B**). In SS-100 iEry however, pomalidomide and decitabine could not increase HbF expression, in line with what we observed for HU and RN-1 (**Fig. 3C**). Treatment with SR-18292^41,42^, on the other hand, resulted in a 1.3-fold increase in HbF-FITC MFI in this patient background that did not respond to several other known HbF inducers, suggesting that peroxisome proliferator-activated receptor gamma coactivator 1-alpha (PGC1α) activation via SR-18292 could represent a viable pharmacological bypass for this highly refractory genetic background. We previously also observed HbF induction by SH6, a small molecule degrader of ZBTB7A, in SS-100 iEry^43^. Together, these findings establish a powerful proof-of-concept for functional drug prioritization. They underscore the platform’s utility to dissect patient-specific drug resistance and identify tailored therapeutic alternatives, ultimately advancing a precision medicine approach for the genetically diverse SCD population.

**Figure 3.**
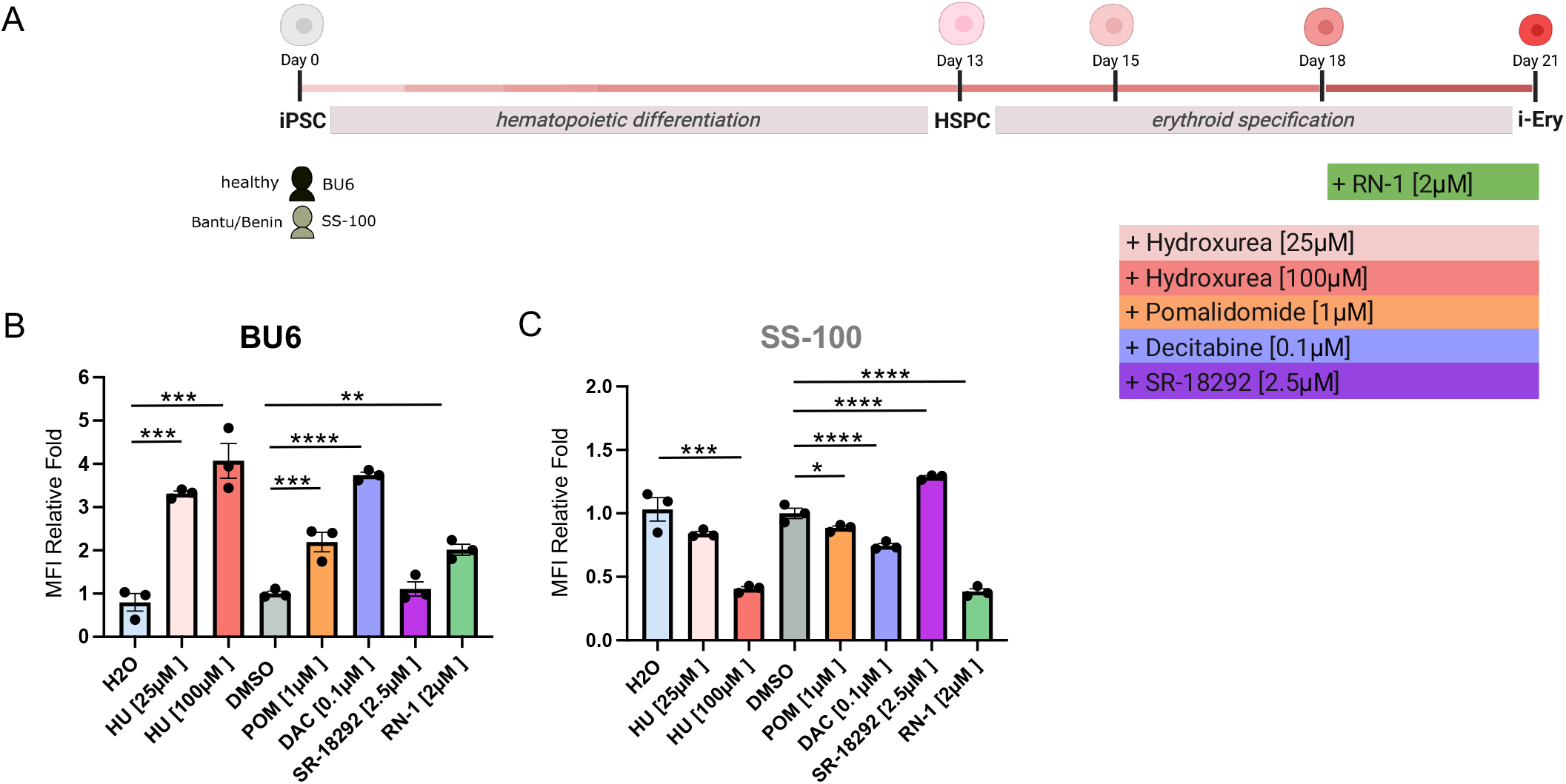
SCD iEry recapitulate patient-specific treatment responses and suggest therapeutic alternatives. A) Schematic overview of differentiation and dosing process. B-C) HbF induction following treatment with different HbF inducing agents, quantified by Mean Fluorescence Intensity (MFI) relative to control condition (n=3) for (B) BU6 and (C) SS-100. Statistical significance between experimental groups was determined using an ordinary one-way ANOVA test with Dunnett correction. A p value <0.05 was considered significant: *p < 0.05, **p < 0.05, ***p < 0.001 and ****p < 0.0001. Error bars represent mean ± standard error of mean (SEM)

### iEry provide a versatile platform to study the impact of drugs on erythroid cell characteristics and differentiation parameters

To evaluate the impact of pharmacological intervention on erythroid maturation, we tracked the expression of Glycophorin A (CD235a) and Band3. We noted that LSD1 inhibitor RN-1 exerts a stage-specific effect on erythropoiesis. When administered shortly after erythroid specification (day 15), RN-1 inhibited the further specification of CD235a^-^ progenitors into the erythroid lineage (**Fig. 4A**). However, cells that had already committed to the erythroid fraction (CD235a^+^) prior to dosing successfully matured from early (Band3^-^) to late (Band3^+^) stages. The block in early erythroid specification upon RN-1 addition was reflected in different proportions of early versus late erythroid fractions in both healthy control and SCD iEry 3 days post dosing (**Fig. 4B**). For BU6, we noted 14.2% (RN-1) versus 77.2% (DMSO) of Band3^-^ cells and the inverse (85.8% versus 22.8% respectively) for Band3^+^ cells at day 18. SA-170 iEry cultures harbored 13% (RN-1) versus 60% (DMSO) of Band3^-^ cells and 87% (RN-1) versus 40% (DMSO) Band3^+^ cells at this time. These findings informed an optimized dosing strategy for RN-1 where we initiated treatment at day 18 to accurately assess HbF induction potential without confounding the results by blocking early erythroid specification.

**Figure 4.**
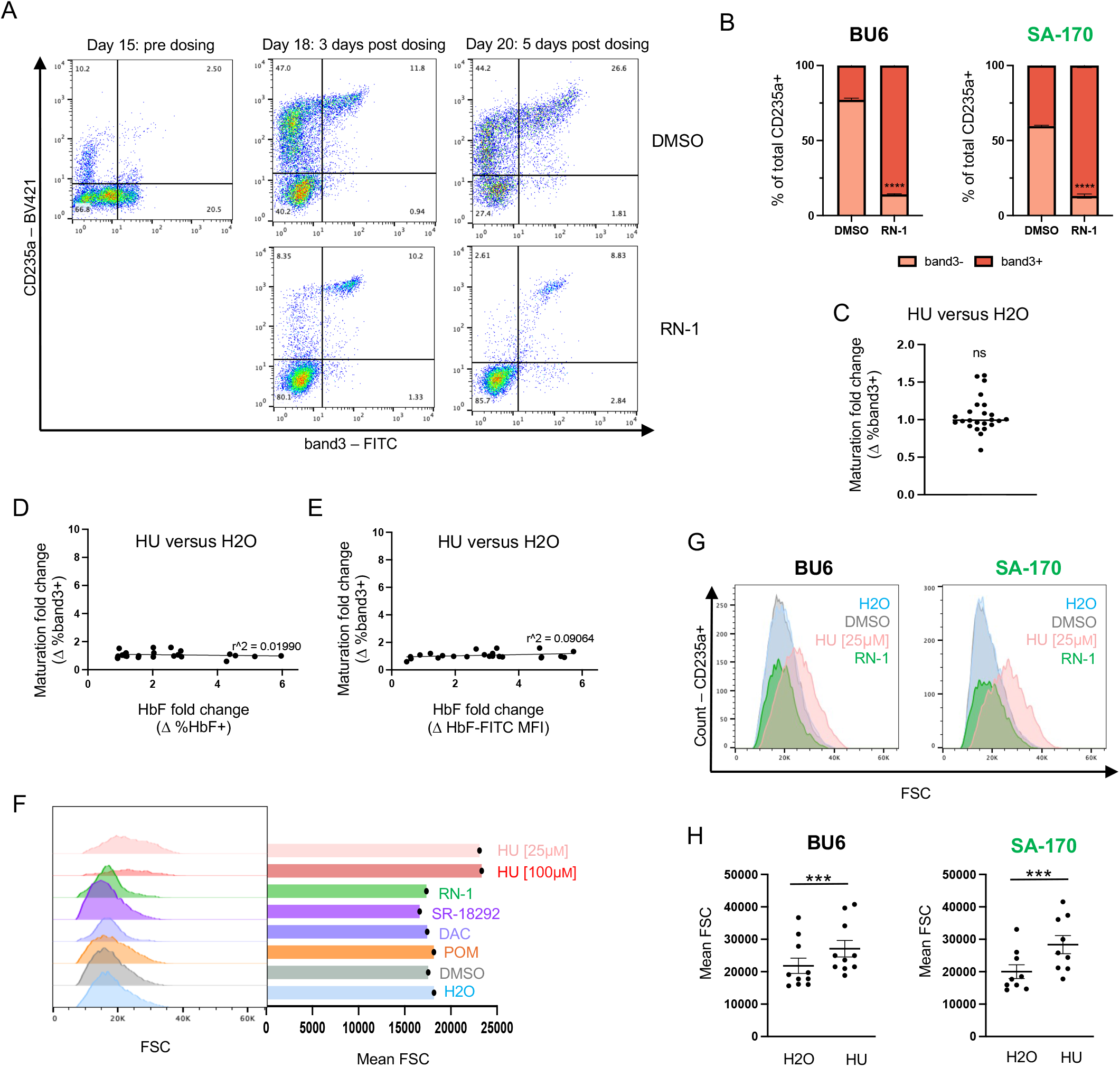
iEry provide a versatile platform to study the impact of drugs on erythroid cell characteristics and differentiation parameters. A) Flow cytometry plots illustrating differentiation progress of cells dosed with RN-1 on day 15 versus the vehicle control (DMSO). B) Quantification of percentage band3- and band3+ cells 3 days post initiation of RN-1 or DSMO dosing withing the CD235a+ fraction (n=3). C) Fold change in maturation (change in %band3+ cells within CD235a+ fraction) between HU and H2O treated conditions following 6 days of treatment for HU responders BU6, SA-170 and SS-24 (n=24). D-E) Fold change in maturation (change in %band3+ cells within CD235a+ fraction) versus fold change in %HbF+ cells (D) or HbF-FITC MFI (E) between HU and H2O treated conditions following 6 days of treatment for HU responders BU6, SA-170 and SS-24 (n=24). F) Comparison of forward scatter (FSC) distribution and mean FSC on day 21, following treatment with HbF inducing agents (BU6). G) Representative histograms illustrating shift in FSC following 6 days of HU treatment in both healthy control (BU6) and SCD (SA-170) iEry. H) Quantification of shift in Mean FSC following 6 days of HU treatment in both healthy control (BU6) and SCD (SA-170) iEry. Statistical significance between experimental groups was determined using a paired Student’s t test (B,H) or a one sample t test (C). A p value <0.05 was considered significant: ns>0.05, ***p < 0.001 and ****p < 0.0001. Error bars represent mean ± standard error of mean (SEM)

We similarly assessed the effect of HU dosing on the maturation process to ensure that the observed HbF induction was not solely due to enhanced maturation of the cells in the presence of HU. We quantified changes in % band3^+^ cells between HU- and H2O-treated iEry from backgrounds that showed HU-mediated HbF induction capacity (BU6, SA-170 and SS-24) and found no difference between treatment and vehicle control, illustrated by a ‘maturation fold change’ of 1.06 (**Fig. 4C**). Meanwhile, corresponding HbF readouts across differentiations did show clear increases in both %HbF expressing cells (**Fig. 4D**) and HbF-FITC MFI (**Fig. 4E**). These results reflect the absence of correlation between changes in HbF expression and maturation (r^2^=0.01990 and r^2^=0.09064 respectively) upon HU treatment.

In addition to drug-specific effects on erythroid differentiation, we also noted drug-specific changes in forward scatter (FSC) of cells as measured by flow cytometry. FSC provides a relative measurement of cell size and as such can be used as a proxy for mean corpuscular volume (MCV), and by extension mean corpuscular hemoglobin (MCH) as the latter two are correlated. Drug-induced changes in MCH could have important implications for the efficacy of a therapeutic agent as cells are thought to be protected from sickling when the ratio of HbF to MCH per cell is ≥1:3^44^. Therefore, compounds that increase MCH might require a larger increase in HbF to achieve this protective ratio. When examining the FSC profiles of cells dosed with different compounds, we noted that HU increased mean FSC (**Fig. 4F-H**), consistent with its reported effect on MCV^45,46^. This effect was unique to HU amongst the drugs tested in our assays and could be observed in both healthy control (BU6) and SCD (SA-170) iEry (**Fig. 4G,H**). Together, these findings illustrate how this SCD-specific renewable cell source enables the dissection of drug-specific effects on cell characteristics and the progression through erythroid differentiation.

## DISCUSSION

While the primary molecular defect in SCD is a shared β-globin point mutation, the clinical manifestation and response to treatment are profoundly shaped by a patient’s broader genetic background^9,18^. Although curative gene therapies offer transformative potential, logistical and financial barriers necessitate the continued development of universally accessible, oral HbF inducers. To de-risk this development pipeline, preclinical screening models must evolve beyond single-genotype systems to accurately reflect the genetic heterogeneity inherent in the patient population.

Our findings establish that patient-specific, iPSC-derived erythroid cells (iEry) provide a high-fidelity platform to capture this biological diversity. Importantly, we show that the inherent baseline expression of HbF in iEry cultures does not obscure the pharmacodynamic induction of HbF by agents such as HU and RN-1, maintaining a dynamic range comparable to primary adult erythroid cultures. Crucially, the platform also captures the *absence* of induction. For some patient backgrounds, HbF induction was never achieved upon HU treatment, even across multiple independent differentiation and dosing experiments and at different drug doses. This outcome mirrors the variable response to HU seen in the SCD patient population^9^. While HU response can be optimized using pharmacokinetically guided dosing^8,47,48^, certain patients fail to meaningfully increase HbF levels, despite long-term adherence^9^. Most notably, the inability of HU to induce HbF in the SS-100 iEry line closely aligns with the refractory clinical history of the donor patient. This correlation strongly suggests that HU resistance extends beyond pharmacokinetic limitations, such as drug absorption or metabolism, and is intrinsically linked to the patient’s specific genetic and epigenetic landscape.

Harnessing this renewable cellular resource, we further explored drug resistance in the SS-100 background and evaluated a panel of canonical HbF inducers spanning distinct mechanistic classes as therapeutic alternatives. In addition to failing to respond to HU, the SS-100 line was refractory to upstream epigenetic modulation (via the LSD1 inhibitor RN-1 and the DNA hypomethylating agent decitabine) as well as IMiD-mediated repressor degradation (pomalidomide). However, treatment with the direct PGC1α activator SR-18292 modestly elevated HbF. We previously also observed that treatment with SH6, a non-IMiD degrader of the independent γ-globin repressor ZBTB7A, successfully achieved HbF induction in SS-100 iEry^43^. This divergent response serves as a powerful proof-of-concept for phenotypic drug triaging. It suggests that while the pathways governing upstream epigenetic derepression and BCL11A-axis modulation may be functionally obstructed in this genetic context, parallel nodes of HbF regulation, specifically ZBTB7A degradation and PGC1α co-activation, remain targetable. By mapping the potential molecular roadblocks in a non-responder’s genetic architecture in vitro, this platform serves as a renewable, functional tool to prioritize the most promising pharmacological bypasses. This capacity to capture and navigate complex patient-specific drug responses streamlines the preclinical development pipeline and represents a vital step toward advancing a precision medicine paradigm in SCD.

Furthermore, clinical efficacy in SCD relies not only on robust HbF induction but also on the preservation of viable red blood cell production. Our platform provides the developmental resolution required to decouple these two processes. For example, we observed that early administration of the LSD1 inhibitor RN-1 disrupts erythroid specification, effectively stalling the transition of CD235a^-^ progenitors into the committed CD235a^+^ compartment. This underscores the delicate balance between targeted epigenetic modulation and normal hematopoietic commitment. Consequently, comprehensive preclinical screening must evaluate both the potency of HbF induction and a compound’s potential footprint on the broader erythroid differentiation trajectory to predict and avoid drug-induced cytopenias. In line with this, we were also able to rule out differences in iEry maturation as the driver of increased HbF expression upon HU treatment.

Additionally, we examined drug-induced changes in FSC as a proxy for MCV and MCH. Compound heterozygotes for HbS-hereditary persistence of fetal hemoglobin (HPFH)^49^ have approximately 10 pg of HbF per red blood cell, an amount sufficient to prevent nearly completely HbS polymerization. The HbF: HbS ratio is ∼0.30, as HbS-HPFH red blood cells typically contain about 26 pg of total hemoglobin^44^. This protective ratio is achieved in all cells due to the pancellular distribution of HbF. Gene therapy approaches similarly result in pancellular HbF reactivation, protecting all edited cells. In contrast, HU induces a heterocellular increase in HbF^44^. HU also increases MCV and MCH. The size of this increase varies, and some cells may not be protected if the increase in MCH outpaces the increase in HbF, reducing the HbF:HbS ratio to < 0.30 and leaving cells susceptible to HbS polymerization and sickling. We show here that these cell characteristics can be modeled and tracked in our iPSC platform, where an increase in FSC was observed upon HU dosing, in line with its reported effect on MCV.

Finally, the intrinsic pluripotency of iPSCs extends the utility of this model beyond hematopoiesis. Because these cells can be differentiated into diverse lineages, this platform offers the unique opportunity to assess the toxicity of novel compounds in patient-matched liver or cardiac tissues, providing a comprehensive, genotype-specific safety profile.

In sum, by capturing the complete genetic background of the individual, this diversity-aware screening approach bridges the gap between monogenic disease modeling and personalized therapeutic discovery. As such, this SCD iPSC platform expands the preclinical toolbox, adding genetic diversity as an important and often overlooked component, ultimately facilitating the development of SCD treatments tailored to the complex biology of the broader patient population.

## AUTHORSHIP CONTRIBUTIONS

B.P. performed experiments, analyzed data, and co-wrote the manuscript; A.G., M.L., M.L.A., A.C., T.T., M.S., and C.W. performed experiments and analyzed data; A.W. analyzed clinical data; M.H.S. and G.J.M. conceived and designed the study and wrote the manuscript; K.V. conceived and designed the study, supervised the research, analyzed data, and wrote the manuscript. All authors reviewed and approved the final version of the manuscript.

## ACKNOWLEDGEMENTS

The authors thank Marianne James from the Center for Regenerative Medicine (CReM) iPSC Core for technical assistance. This project was supported by the National Center for Advancing Translational Sciences, National Institutes of Health, through BU-CTSI Grant Number 1UL1TR001430, awarded to KV. Its contents are solely the responsibility of the authors and do not necessarily represent the official views of the NIH. This work was also supported through an Early Career Scientific Research Grant from the Association for the Advancement of Blood & Biotherapies (AABB, formerly known as National Blood Foundation) for KV and a collaborative grant between the Murphy and Steinberg Groups (1R01HL133350) from the National Heart, Lung, and Blood Institute, National Institutes of Health.

---

**Figure S1.**
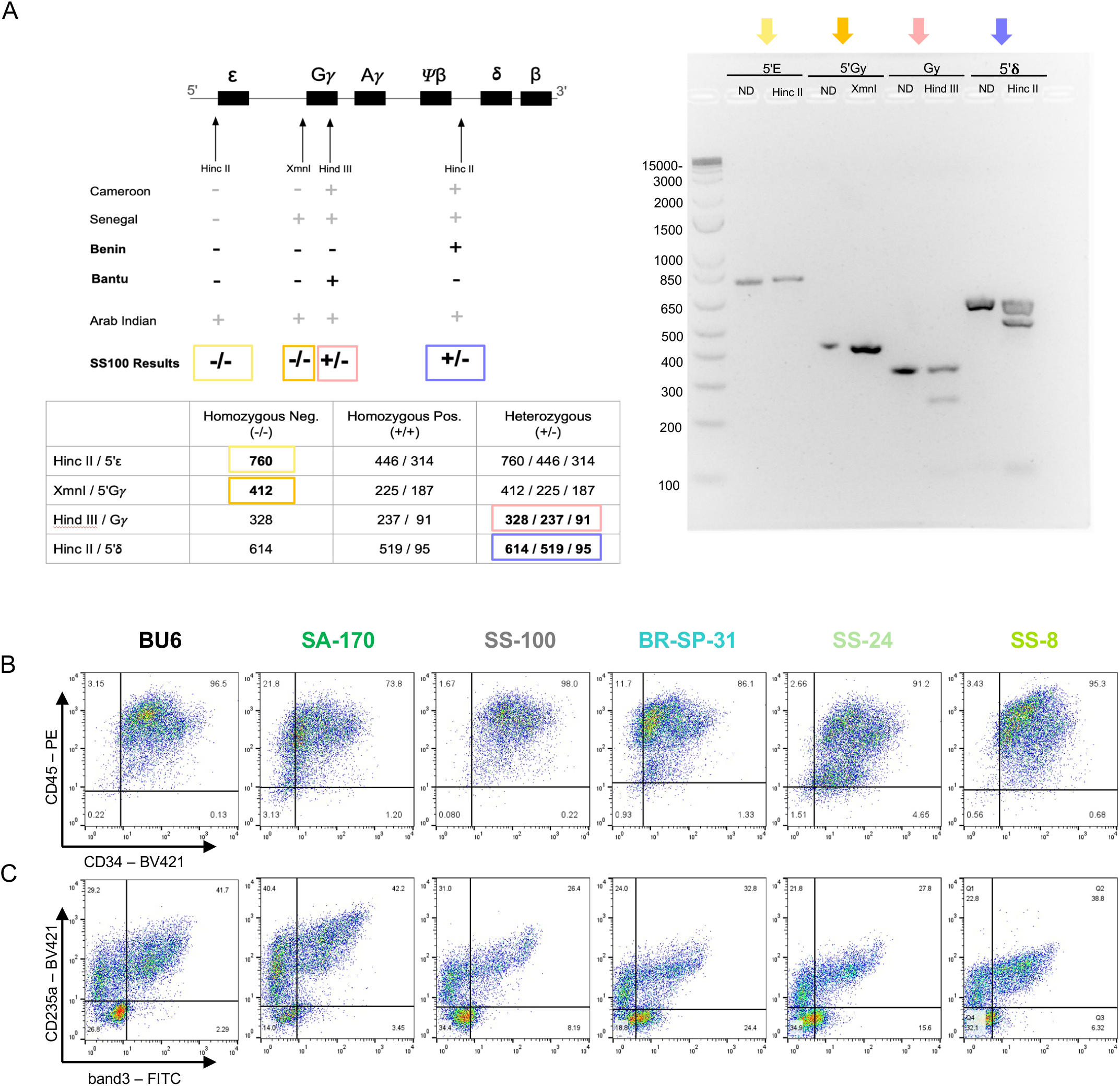
Haplotyping and differentiation progression. (A) Haplotyping of SS-100 iPSCs (benin/bantu) via restriction fragment length polymorphism (RFLP) assay. ND = non-digested control B-C) Flow cytometry plots illustrating differentiation progress of cells corresponding to different backgrounds on (B) day 13 and (C) day 21.

**Table S1.**
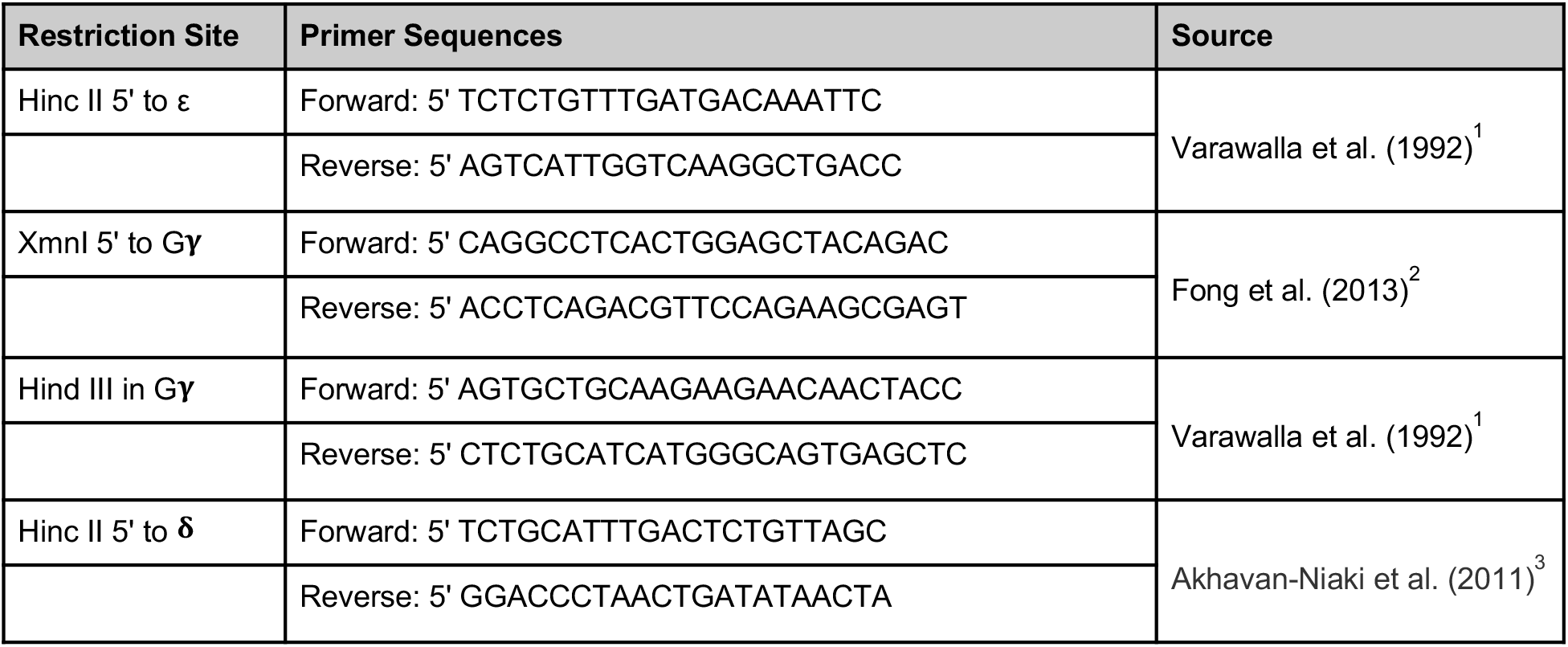
Haplotyping primers. Sequences and references indicated^1-3^

